# Quantifying Asymmetric Coevolutionary Dynamics using Normalized Phylogenetic Costs

**DOI:** 10.64898/2026.06.29.734822

**Authors:** Sanket Wagle, Alexey Markin, Tyler J. Sherman, Christie Mayo, Tillie J. Dunham, Corey Brelsfoard, Lee W. Cohnstaedt, William C. Wilson, Tavis K. Anderson, Oliver Eulenstein

## Abstract

Coevolutionary studies aim to characterize associations, such as virus-host relationships, by using phylogenetic distances to quantify the topological concordance between the phylogenies of interacting taxa. However, phylogenetic distances cannot capture asymmetrical relationships that arise from differences in sampling, evolutionary rates, or characterizations between datasets. Furthermore, a lack of accurate normalization complicates the interpretation and validation of coevolutionary analyses. To address these limitations, we employed the Asymmetric Cluster Affinity and Cluster Support costs as a general framework to quantify coevolutionary patterns across multiple biological scales. We benchmarked the precision of these costs by reanalyzing a curated dataset documenting interspecies transmission frequencies across nineteen virus-host phylogenies. Our results corroborate prior findings showing that all virus families under study can cross species boundaries; however, the asymmetric costs provide a more granular representation, demonstrating that the frequency of such events varies significantly across families. We then applied the Asymmetric Cluster Support cost to quantify preferential gene segment pairings within the Bluetongue virus genome. This analysis revealed a close phylogenetic association between the outer capsid proteins VP2 and VP5, likely reflecting shared selective pressures due to their critical roles in cell entry and exit. In contrast, gene segments encoding nonstructural proteins exhibited discordant evolutionary histories relative to other segments. Finally, we demonstrated that the Asymmetric Cluster Support cost can detect coevolutionary dynamics in swine influenza A virus, identifying novel gene pairings indicative of major viral reassortment events. Overall, our approach demonstrates that normalized asymmetric phylogenetic costs accurately capture complex biological relationships and provide a robust framework for quantifying fine-scale coevolutionary dynamics in rapidly evolving pathogens.

## Introduction

Coevolution occurs when two or more groups of taxa affect each other’s evolutionary history [Thompson and Pagel, 2001] in the form of beneficial or deleterious interactions such as mutualism and parasitism [Anderson and Sukhdeo, 2011, Dismukes et al., 2022]. In the context of hosts and their pathogens, some pathogens infect and have coevolved within a single host species [Drezen et al., 2017, Harrach et al., 2019, Zinner et al., 2023], whereas others have a broader host range [Neumann et al., 2009, Duque-Valencia et al., 2019]. The history of these associations can be detected and quantified through the degree of topological congruence between the host and pathogen phylogenies [Avino et al., 2019a, Geoghegan et al., 2017]. In addition, major evolutionary changes such as interspecies transmissions where a pathogen jumps to a new host species [Olival et al., 2017], and virus transmission dynamics within and between hosts may be observed within the topological structure of phylogenies [Paterson et al., 2010, Carius et al., 2001]. While coevolutionary studies traditionally focus on comparing distinct species or families, analogous patterns occur at the genomic scale, where phenomena such as preferential gene segment pairings [Zeller et al., 2021, Neverov et al., 2015] can be characterized as tightly linked coevolutionary processes. Ultimately, quantifying such phenomena across both interspecies and intra-genomic scales reduces to the same underlying mathematical problem: accurately measuring the topological discordance between phylogenies.

Detecting coevolutionary patterns is often achieved by constructing a “tanglegram” to visualize host-pathogen associations across multiple phylogenies [Page, 2003]. Topological congruence between the phylogenies suggests coevolution between genes, or potentially a pattern of co-speciation between hosts and their pathogens [Morand et al., 2000, Brooks, 1985, Paterson et al., 1997, Machado et al., 2005]. Conversely, a tanglegram with crossing edges and topological incongruence suggests divergence and can serve as an indicator of new gene pairings acquired through reassortment, or discordant evolutionary dynamics. However, the tanglegram approach that has been widely used in virus biology [Page, 2003, Castelán-Sánchez and Poon, 2025] has limitations when applied to coevolutionary studies; the entanglement can be explained by a large number of different evolutionary events [Bininda-Emonds and RP, 2004] and estimating the optimal number of events for a tanglegram becomes exceedingly difficult as the number of species and taxa increase [Ovadia et al., 2011]. Moreover, tanglegrams are qualitative tools and thus unsuitable for large-scale quantitative analysis of coevolutionary processes.

Hence, to overcome the limitations of tanglegrams, phylogenetic distances have been used as an alternative measure of coevolutionary divergence [Geoghegan et al., 2017, Avino et al., 2019a, Jones et al., 2021, Brooks et al., 2016]. These distances begin by converting phylogenies into equivalent representations, such as clusters [Lin et al., 2011, Bogdanowicz and Giaro, 2013, Robinson and Foulds, 1981], subtrees [Estabrook et al., 1985], or kernel mappings [Poon et al., 2013], and then mapping these representations onto a metric space. The evolutionary divergence between phylogenies is then quantified as the distance between them in this metric space. This method allows phylogenetic distances to possess mathematically favorable properties, such as symmetry, and has led to their application in a variety of phylogenetic analyses [Chaudhary et al., 2013, Giardina et al., 2017, Bogdanowicz and Giaro, 2017].

However, relationships between phylogenies are often fundamentally asymmetric. This asymmetry can be caused by systemic factors such as sampling bias [Bush et al., 2000, Troudet et al., 2017] and a lack of phylogenomic data [Rinke et al., 2013] or by biological phenomena, such as different evolutionary rates and selective pressures [Zeller et al., 2021, Erasmus and Huismans, 1981]. Relying on phylogenetic distances to analyze such phylogenies forces a symmetric assumption that ignores these asymmetric relationships and can thus yield misleading results. Phylogenetic distances are further hindered by a lack of accurate normalization. Since distance values are often affected by factors such as taxon size, topological distributions, and structural constraints [Smith, 2021], identical numerical values can represent fundamentally different levels of discordance across datasets. Without normalizing these values, it is impossible to differentiate between meaningful biological variance and artifacts of topological structure. Unfortunately, most phylogenetic distances only provide asymptotic bounds [Lin et al., 2011, Bogdanowicz and Giaro, 2017] and thus cannot be adequately normalized, leading to biases that prevent comparisons within and between studies. Hence, an ideal approach to detect coevolution should be able to represent the inherent asymmetry between the phylogenies as well as enable accurate and efficient normalization based on the input topologies.

In this work, we investigate the utility of two asymmetric phylogenetic costs, the normalized Asymmetric Cluster Affinity (ACA) and Cluster Support (ACS) costs [Wagle et al., 2024] in identifying coevolutionary patterns. By relaxing the requirement for symmetry, these costs can accommodate the directional nature of coevolutionary relationships and offer complementary biological insights when comparing phylogenies: the ACA cost measures raw differences between clusters, placing greater emphasis on disagreements involving larger clades, whereas the ACS cost evaluates differences as a percentage of cluster size, making it sensitive to localized topological shifts. To demonstrate the broad utility of this approach, we apply the ACA and ACS costs across three distinct biological case studies: (1) validating their performance on an empirical virus-host cophylogeny dataset based on analyses by Geoghegan et al. [Geoghegan et al., 2017]; (2) detecting stable gene pairings within whole-genome sequences of the Bluetongue virus; and (3) identifying preferential gene pairings and novel reassortment in influenza A viruses in swine. Our analysis demonstrates how normalized asymmetric costs provide a versatile framework capable of quantifying coevolutionary patterns across multiple evolutionary scales.

## Materials and Methods

### Basic definitions and terminology

A *phylogenetic tree T* over a taxon set *M* is a rooted tree where all the edges are pointed away from its root, and its leaves are identified (i.e., bijectively labeled) with the elements from *M*. The set of vertices and edges in *T* are denoted by *V* (*T*) and *E*(*T*), respectively. For a tree *T*, we define the *leaf labeling* of *T* as the bijection Λ*_T_* : *L*(*T*) → *M*. For a node *v* in *V* (*T*), we define the *cluster* of *v* as the set of all leaves descendant from *v* and denote it by *c_v_*. A *cluster set* of *T*, denoted by *C*(*T*), is the set of all clusters associated with the nodes in *T* and uniquely identifies the tree *T* [Semple and Steel, 2003]. These cluster sets form the basis of the widely used Robinson-Foulds(RF) distance [Robinson and Foulds, 1981], which defines the topological distance between two trees as the symmetric difference between their cluster sets.

The RF distance, however, is extremely sensitive to minor topological differences [Lin et al., 2011] and assumes a symmetric relationship between the trees being studied, rendering it less suitable for coevolutionary studies involving considerable noise or asymmetric relationships. To enable such asymmetric comparisons, the ACA and ACS costs were introduced as extensions of the RF distance. Formally, the ACA cost from a cluster *c* to a tree *T* is defined as *d*(*c, T*) := min*_x_*_∈_*_C_*_(_*_T_* _)_ |*c*Δ*x*| while the ACS cost from the same cluster to the tree *T* is defined as 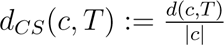. Similarly, for trees *S, T*, the ACA cost from *S* to *T* is defined as *d*(*S, T*) := Σ*_c_*_∈_*_C_*_(_*_S_*_)_ *d*(*c, T*) and the ACS cost from *S* to *T* is defined as *d_CS_*(*S, T*) := Σ*_c_*_∈_*_C_*_(_*_S_*_)_ *d_CS_*(*c, T*).

Unlike the RF distance, which evaluates the clusters as indivisible units, the ACA and ACS costs allow partial matches and evaluate the information within each cluster pair. This approach allows the ACA and ACS costs to capture more nuanced tree differences and be resilient to small topological errors [Wagle et al., 2024]. For example, consider trees *T*_1_ and *T*_2_ in Fig 1. While *T*_1_ and *T*_2_ differ only in the placement of the {*a*} taxon, the RF distance characterizes both trees as being maximally different and equivalent to the case where every single taxon was misplaced. Conversely, the ACA cost from *T*_1_ to *T*_2_ is slightly more than half of the maximum ACA cost for *T*_1_, correctly indicating that while the shift of the *a* taxon is significant, the two trees are still broadly similar.

**Fig. 1.**
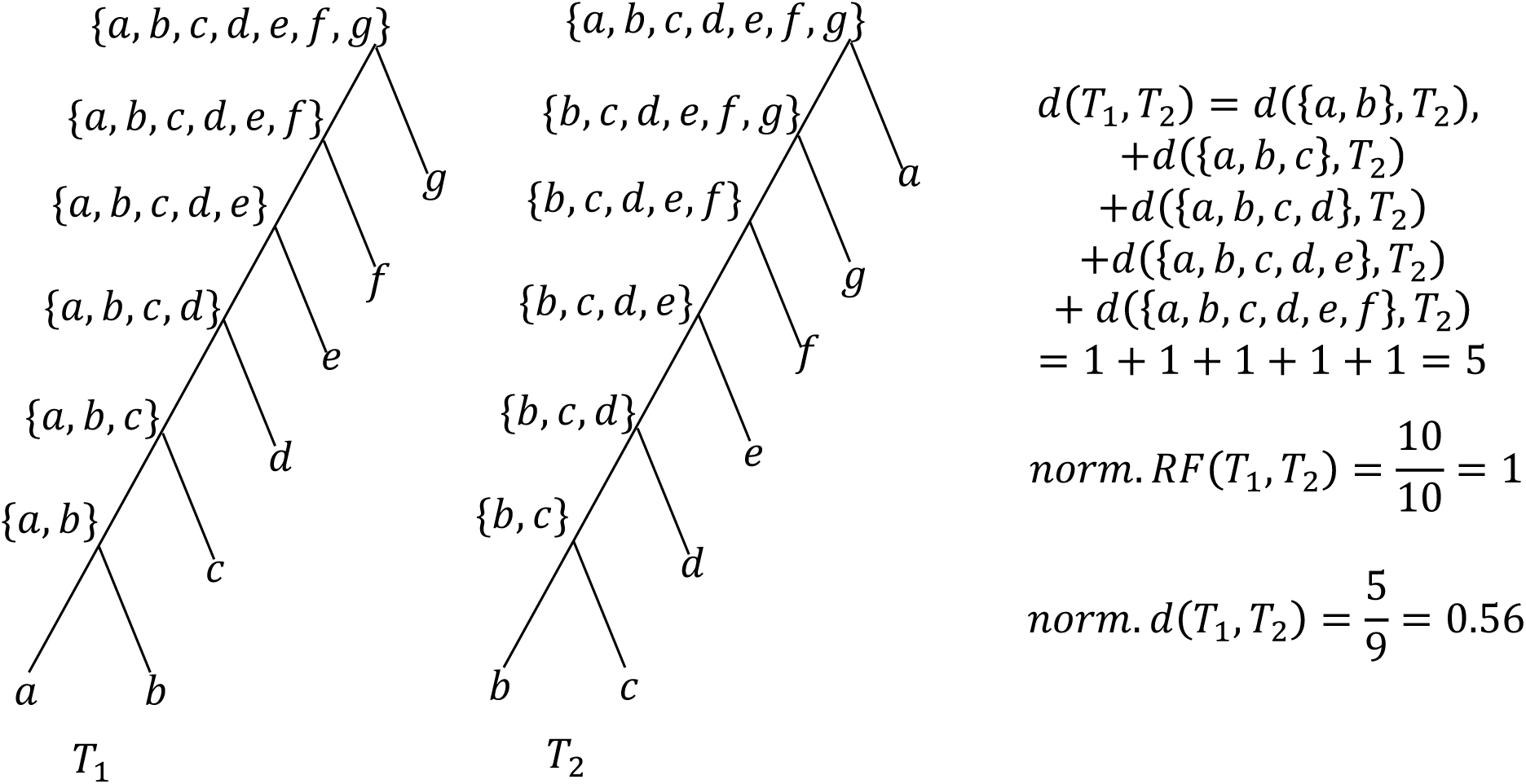
Comparison of Robinson-Foulds distance and Asymmetric Cluster Affinity cost. The Robinson-Foulds distance considers the clusters as indivisible units and considers the trees as being maximally different. On the other hand, the ACA cost considers the information within the clusters and assigns a normalized cost of 0.56 to map *T*_1_ to *T*_2_. For brevity, we ignore the root and leaf clusters when computing the ACA cost since they are present in both trees.

### Normalized costs

Normalizing a phylogenetic cost requires analytical solutions for the maximum value of the cost for two fixed input topologies, referred to as the diameter of the cost. Formally, the diameter of a phylogenetic cost *f* for trees *S, T* is defined as the maximum value of *f* over all possible leaf labelings of *T* and is denoted as *diam*(*f, S, T*). Hence, given fixed leaf labelings for *S* and *T*, we define the normalized version of the cost *f* as 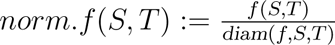. Here, we describe the diameters for the RF distance and ACA costs given two input topologies and derive the diameter for the ACS cost.

Lemma 2.1 (From [Robinson and Foulds, 1981]) For a pair of rooted trees *S, T* over the leaf set *M*, *RF* (*S, T*) ⩽ 2|*M* | − 4.

Based on a previous result by Wagle et al [Wagle et al., 2024] we can derive the maximum bounds on the ACA and ACS costs between two trees *S, T* as follows

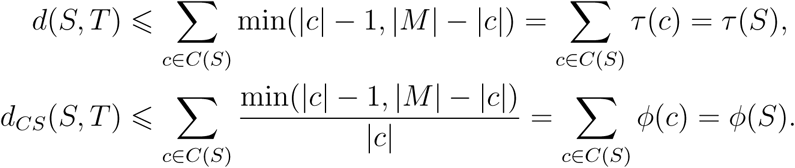

Lemma 2.2 (From [Górecki et al., 2026], Theorem 3, ACA Diameter for Fixed First Topology) For a pair of trees *S, T*, there exists a leaf labeling Λ*_T_* such that *d*(*S, T*) = *τ* (*S*).

Lemma 2.3 (From [Wagle et al., 2024], Lemma 8) For any two trees *S, T*, *d_CS_*(*S, T*) ⩽ *ϕ*(*S*).

Corollary 2.4 For a pair of trees *S, T*, there exists a leaf labeling Λ*_T_*such that *d_CS_*(*S, T*) = *ϕ*(*S*).

*Proof.* Let *c* be a cluster such that *c* ∈ *C*(*S*). Note that if *d*(*S, T*) = *τ* (*S*), then *d*(*c, T*) = *τ* (*c*). It follows that 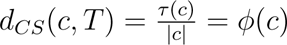 and *d_CS_*(*S, T*) = *ϕ*(*S*). Hence, utilizing the leaf labeling described in Theorem 2.2, we get *d*(*S, T*) = *τ* (*S*) which implies *d_CS_*(*S, T*) = *ϕ*(*S*).

### Virus-host phylogenies dataset collection

To analyze the performance of the ACA and ACS cost in quantifying coevolutionary patterns, we recreated a study by Geoghegan et al [Geoghegan et al., 2017] that quantified the frequency of interspecies transmission for virus families and their respective host species. We obtained 19 virus and host phylogenies from a Github repository from the original study [Geoghegan and Duchêne, 2017]. In order to facilitate comparisons between our study and the original results, we maintained the taxonomy used in the previous analysis; however, some of these family designations have been revised (e.g. Reoviridae is now a taxonomic order, not a family [Walker et al., 2022]). The trees used in the original analysis were unrooted, and as ACA and ACS are rooted tree costs, we used midpoint rooting to infer a root for all trees. The choice of direction for the ACA and ACS costs was determined by the significantly larger number of polytomies in the host phylogenies than the virus phylogenies; this difference was a combination of host phylogenies having lower phylogenetic signal and multiple virus species infecting the same host. Under a model of no interspecies transmission, due to the difference in polytomies, we would expect the host phylogenies to completely contain the virus phylogeny, i.e., the clusters in the host phylogenies would either be present in the virus phylogenies or be supersets of the clusters in the virus phylogenies. In the context of the ACA and ACS costs, the cost scores from the host phylogeny to the virus phylogeny under a model of no interspecies transmissions and no error would be zero, whereas for the same phylogenies in the reverse direction, the difference in polytomies would inflate the cost scores erroneously indicating divergence. Hence, to avoid the bias induced by the difference in polytomies, we computed the ACA and ACS costs from the host to the virus phylogenies and calculated the rooted Robinson-Foulds (RF) distance between the phylogenies. Note that the direction of computation is irrelevant for the RF distance since it is symmetric. However, this symmetric assumption also leads to inflated values as the RF distance assumes that polytomies are indicators of evolutionary divergence rather a difference in phylogenetic resolution.

### Detecting coevolutionary patterns between Bluetongue virus genes

Bluetongue virus (BTV) belongs to the Orbivirus genus within the order Reovirales and is responsible for Bluetongue disease in ruminants, causing severe economic damage either due to the direct effect on livestock or indirectly via trade restrictions and the cost of surveillance and control [Mellor and Wittmann, 2002, Mellor et al., 2008]. To quantify and characterize the coevolutionary relationships between the segments of BTV, we performed a pairwise comparison between the gene trees for each segment using the ACS cost. We inferred gene trees using IQ-Tree v2.0.7 [Minh et al., 2020] for each segment using published sequences from NCBI Virus [Brister et al., 2015] and new sequence data held within the private client database of the Colorado State University Diagnostic Medicine Center while considering *EHDV-1* as an outgroup. For each pair of segments (*a, b*), we calculated the ACS cost from the segment tree *T_a_* to *T_b_* and vice-versa and visualized the results as a heatmap matrix (Figure 3). As the ACS cost does not impose any structural constraints on the two trees other than the rooting state, we did not resolve any polytomies.

### Influenza A virus dataset collection and curation

To test whether ACS cost scores could detect coevolution of IAV surface proteins, the strength of the association, and if this was correlated with duration of circulation in pigs, we collated publicly available genetic sequence data fetched from octoFLUdb (https://github.com/flu-crew/octofludb). The HA and NA genes were aligned using mafft v7.526 [Katoh and Standley, 2013], the HA clade identity was assigned using the octoFLU pipeline [Chang et al., 2019], and the HA and NA gene trees were sampled into their respective HA clades using smot v0.17.4 [Arendsee et al., 2022]. To minimize the impact of sampling biases present in genomic surveillance data derived from passive surveillance [Janzen et al., 2025], each HA IAV clade and its paired NA gene were randomly sampled to 50 taxa chosen uniformly, and the ACS cost from the HA tree to the NA tree was reported. The uniform random sampling was replicated 100 times for each clade, and the results were summarized (Figure 4).

## Results

### Coevolutionary patterns between virus-host phylogenies

Over the previous century, multiple pandemics have occurred following the transmission of an animal pathogen to humans [Jones et al., 2008]. Hence, viruses circulating in animals are not only a threat to animal health, but may also represent a potential threat to humans [Carroll et al., 2018]. Some examples include influenza A viruses that emerged in swine and resulted in the first pandemic of the 21st century [Garten et al., 2009] and the SARS-CoV-2 pandemic [Worobey et al., 2022, Pekar et al., 2022]. These host switching events significantly alter the evolutionary history of a virus and are key to understanding the central mechanics of viral evolution [Olival et al., 2017, Emerman and Malik, 2010]. However, while interspecies transmission events may have a significant impact, determining the relative frequency and identifying which animal pathogens or viral families cross species boundaries is challenging [Olival et al., 2017, Kaján et al., 2020].

Recently, a study quantified the frequency of interspecies transmission for 19 virus families and their respective animal and human hosts [Geoghegan et al., 2017]. The study aimed to identify interspecies transmission rates by using a normalized version of the unrooted Robinson Foulds (RF) distance (the nPH85 distance) to quantify coevolution between phylogenies comprising of viruses and their respective hosts. While the study was able to detect the ubiquitous presence of interspecies transmission events across all of the studied viral families, it was unable to detect a strong signal for variation between different families. A potential reason for this is the sensitivity of the RF distance to small changes in tree topology, with a phylogenetically detected single interspecies transmission event being indistinguishable from multiple interspecies transmission events within the same group.

Furthermore, the symmetric approach used by the RF distance is unable to account for the difference in the number of polytomies between the host and virus phylogenies. Instead, the RF distance assigns a higher value to such pairs of phylogenies, as the more clearly defined clusters in the virus phylogeny are seen as evidence of topological discord. Hence, while the RF distance performed well as an indicator, it was not able to quantify the relative degree of interspecies transmission across virus families. We hypothesized that the asymmetric nature and more intensive approach used by the ACA and ACS costs would allow a more accurate assessment of the frequency of interspecies transmission events and be able to characterize the variation in interspecies transmission events across virus families.

We observed differences between the results reported in the Geoghegan et al. [Geoghegan et al., 2017] study and the ACA and ACS costs (Figure 2). The ACA and ACS cost values exhibited significantly higher variation across the virus families compared to the RF distance. Specifically, the RF distance had a standard deviation of 0.082 while the ACA and ACS costs had a standard deviation of 0.136 and 0.128 respectively. This increase in variation allowed for a more granular representation of the variation in host-switching events across the virus families. For example, while the *Rhabdoviridae* and *Reoviridae* families exhibited a similar RF distance, *Rhabdoviridae* has a significantly lower ACA cost compared to *Reoviridae*, implying that viruses in the *Reoviridae* family exhibit host switching much more frequently as opposed to viruses in the *Rhabdoviridae* family. *Orthomyxoviridae* was characterized by the ACA cost as having the highest risk for interspecies transmission. Our analyses also demonstrate that *Adenoviridae* may not frequently jump host species and potentially have a lower pandemic risk supporting some empirical characterization studies [Harrach et al., 2019]. We hypothesize that the inflated values in the prior analyses were a result of the Adenoviridae host phylogeny containing a significant number of polytomies representing multiple viruses infecting the same host species; a feature that was absent in the virus phylogeny. An important outcome of our analyses is the wide range of ACA and ACS costs generated, supporting prior studies that show a wide variation in interspecies transmission between virus families [Olival et al., 2017, Emerman and Malik, 2010].

**Fig. 2.**
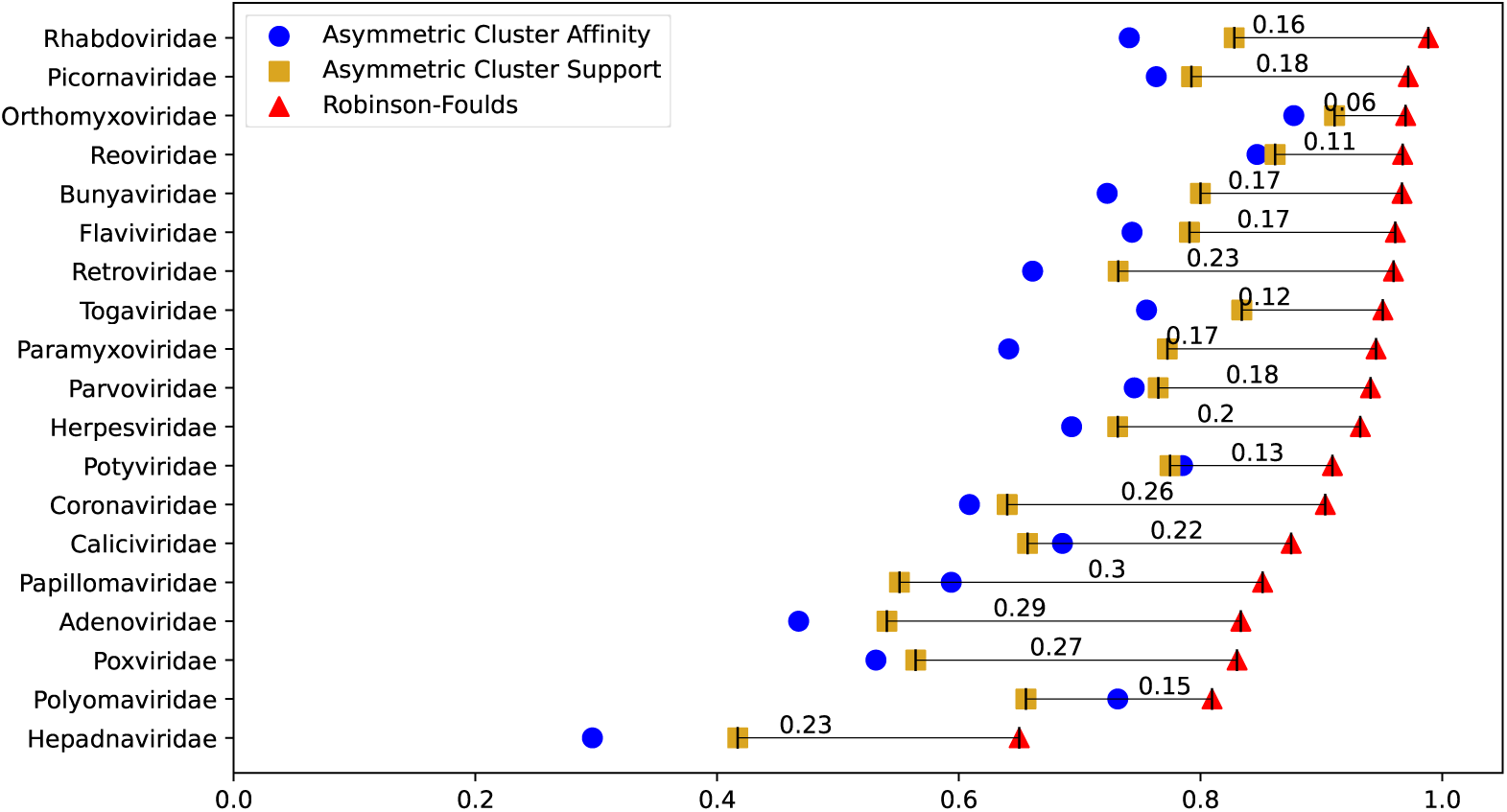
The Robinson Foulds distance and the Asymmetric Cluster Affinity and Cluster Support costs from the host phylogenies to the virus phylogenies. The x-axis shows the relative cost between the phylogenies normalized on a scale from 0 to 1, while the y-axis lists the different virus families in order of their ascending RF distance.

### Coevolutionary patterns between Bluetongue virus genes

Understanding the dynamics of inter-gene linkage can inform and help understand the evolution of RNA viruses [Hufnagel et al., 2023, Zeller et al., 2021]. In addition to rates of evolution of individual gene segments and the impact of these inherent rates on genetic changes observed in circulating viruses, the observed fitness of viruses likely depends on the coordinated function of combinations of genes. Indeed, the function of one gene likely affects the function of other genes and overall viral fitness [Wagner et al., 2000, Hufnagel et al., 2023, Zeller et al., 2021]. Thus, developing novel analytical measures that can identify when there is coordinated evolution between particular proteins can provide insight into whether novel combinations may have a fitness benefit [Thomas et al., 2023].

Bluetongue virus (BTV) is the prototype virus of the ruminant- and camelid-infecting *Oribivirus* genus within the Reovirales order and comprises ten linear segments of double-stranded RNA (dsRNA). The BTV segments encode seven structural proteins and five non-structural proteins involved in viral replication in arthropod and mammalian hosts [Patel and Roy, 2014]. Of these proteins, the VP2 and VP5 that are encoded by segments 2 and 6 collectively form the outer capsid and represent the primary antigenic determinant for the virus [Roy, 2017]. Prior analyses have suggested that strong selection pressure on VP2 and VP5 has resulted in higher sequence variation in Seg2 and Seg6 relative to other gene segments, and consequently, Seg2 is used as the primary identifier for assigning the viral serotype [Erasmus and Huismans, 1981]. The genome segments of BTV have been suggested to evolve independently of one another while replicating in a vertebrate host or arthropod vector [Nomikou et al., 2015]. However, essential interactions on the biochemical level should result in coevolutionary relationships between viral proteins and nucleic acids that can be detected in phylogenetic tree topology.

We observed that the Seg2 and Seg6 phylogenies showed the lowest ACS cost between each other, while exhibiting a relatively higher cost to the other gene segment phylogenies (Fig 3). This suggested a relatively tight coevolutionary history, indicative of their critical role in the formation of the outer capsid [Forzan et al., 2007, Roy, 2017], and also indicated divergence from the other gene segments as a result of the high sequence variation exhibited. We also noted that the Seg10 gene segment phylogeny had, on average, relatively high ACS costs mapping to and from all other gene segment phylogenies. This may reflect a relatively faster evolutionary rate for the Seg10 gene [Alkhamis et al., 2020] as well as stronger selective pressures due to a functional role associated with mediating virulence [Rodríguez-Martín et al., 2021, Han and Harty, 2004].

**Fig. 3.**
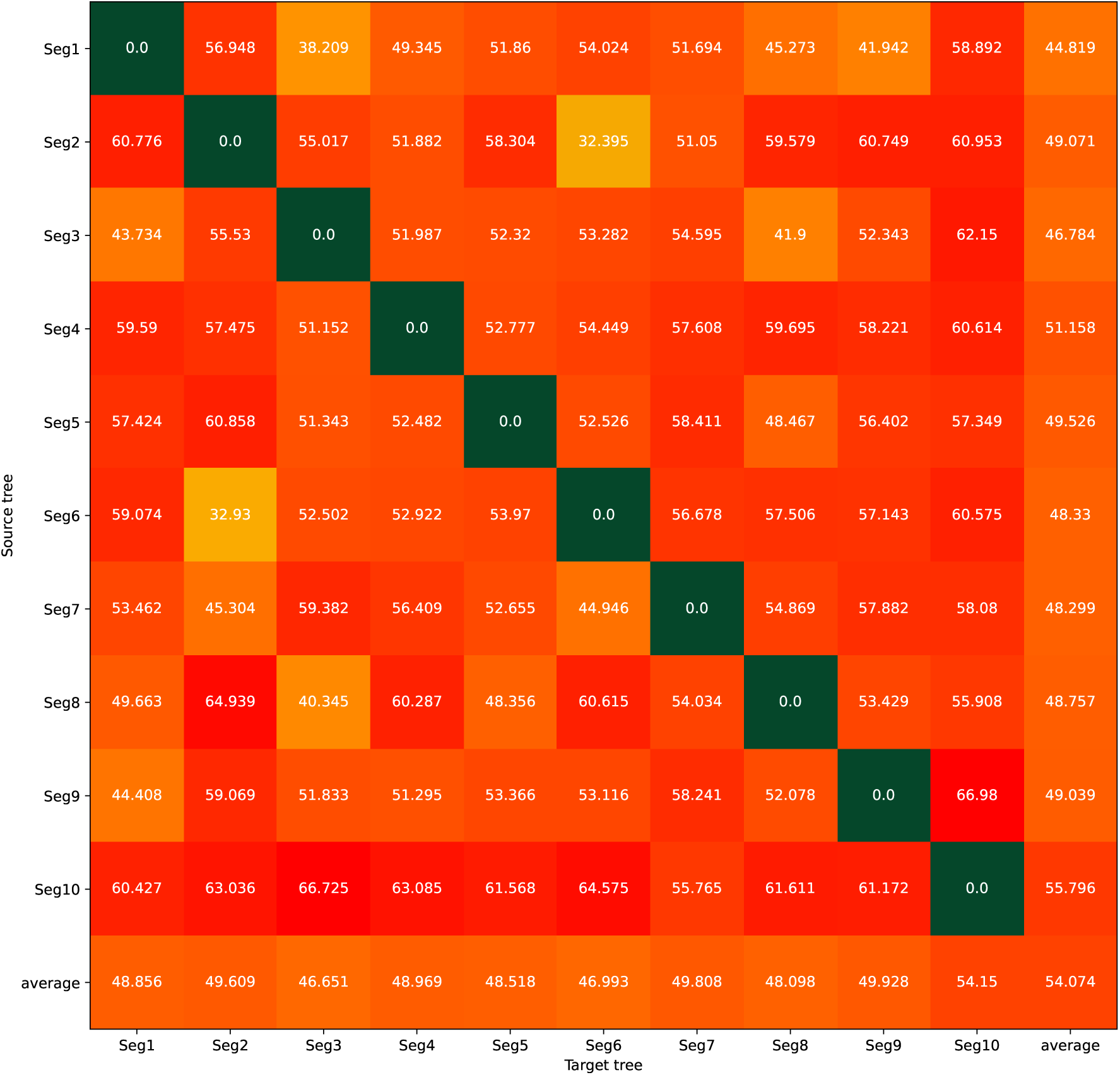
Heatmap of the ACS cost between BTV gene segment phylogenies. Each square represents the ACS cost between a unique source and target pair, with the source phylogeny being specified by the row and the target phylogeny being specified by the column. The last column and row summarize the average ACS cost observed along their respective axes. All values are given as a percentage of the diameter, i.e., the maximum ACS cost achievable by the source phylogeny.

### Preferential gene pairings and reassortment in the surface protein genes of influenza A virus in swine

All endemic H1 and H3 subtype influenza A viruses (IAV) circulating in swine in the United States are derived from human-to-swine transmission events [Anderson et al., 2021]. These viruses are members of the Orthomyxoviridae family, which has shown evidence of frequent interspecies tranmission events (Fig 2). The H1 subtype includes viruses classified as 1A classical swine lineage that emerged coincident with the 1918 human influenza pandemic. Over the last century, the 1A viruses have evolved regionally in the US [Anderson et al., 2021, Walia et al., 2019, Janzen et al., 2025]; and occasionally, novel 1A viruses are introduced as pigs move between countries (e.g., [Nelson et al., 2017]). This has resulted in endemic viruses that share an evolutionary history with the 1918 influenza pandemic, i.e., 1A.3.3.3, or viruses that share a common ancestor with the 1918 pandemic but were introduced into US swine herds more recently, i.e., the 1A.1.1.3 group was introduced into the United States from Canada in approximately 2015 [Nelson et al., 2017]. A second H1 subtype lineage is regularly detected in US pigs, and these viruses were the result of two separate human-to-swine spillover events associated with human seasonal viruses circulating in the early 2000s [Nelson et al., 2011, Vincent et al., 2009]. These viruses have evolved over the past 20 years into three phenotypically distinct groups that are classified as 1B.2.1, 1B.2.2.1, and 1B.2.2.2 [Vincent et al., 2009, Nelson et al., 2011, Rajao et al., 2018]. The H3 subtype IAV reflects a similar tangled evolutionary history with human seasonal IAV, and there have been three independent human-to-swine transmission events [Arendsee et al., 2021]. The first H3 lineage was introduced in the 1990s (1990.4 lineage), with the two other endemic lineages detected in the 2010s (2010.1 and 2010.2 lineages) [Walia et al., 2019, Rajão et al., 2015, Sharma et al., 2022].

Each of the endemic H1 and H3 IAV in pigs is classified using the diversity of the hemagglutinin (HA) surface protein, but the replication of IAV within, and transmission between, hosts is the result of coordinated function between the HA and the other major surface protein, neuraminidase (NA) [Escalera-Zamudio et al., 2020, Neverov et al., 2015]. Consequently, we hypothesized that the HA and NA IAV genes detected in US swine would exhibit preferential gene pairings and, due to their biological function, the strength of the pairing would reflect the duration of circulation in pigs.

There was a trend between the duration of circulation in swine and the coevolutionary pairing strength between IAV proteins measured using the ACS cost (Fig 4). This was most apparent for the HA clade that had been circulating in swine for the longest time, i.e., the 1A.3.3.3 classical swine lineage HA-NA pairing has been in pigs since the 1918 human influenza pandemic. The 1A.3.3.3 scoring suggested a stable HA-NA coevolutionary pairing indicated by a low ACS cost. This result mirrors prior analyses that argued the 1A.3.3.3 HA that was paired with an N1 NA gene had reached a fitness peak [Hufnagel et al., 2023]. Conversely, clades that were more recently introduced into US swine had higher cluster support costs, with a weak trend towards higher ACS costs for the more recent introductions into US pigs. The H3.2010.2 lineage was identified in 2016, and is the most recent human-to-swine spillover that has persisted in US swine. An exception to this pattern was observed in the 1A.1.1.3 clade, which showed a higher ACS cost when compared to viruses that were circulating in pigs for approximately similar times, e.g., H3.2010.1 was scored similarly to the H1 1A.1.1.3. In this example, a reassortment event where the HA acquired new NA gene pairings occurred in approximately 2020 [Janzen et al., 2025]. To investigate this, we split the 1A.1.1.3 phylogeny based on date of sequence collection into pre-2020 and post-2020 phylogenies. We demonstrated that the post-2020 phylogenies have a much higher ACS cost than their pre-2020 counterparts, indicating novel gene pairings due to recent reassortment. These data suggest that the ACS cost may be applied to identifying novel genetic pairings acquired through reassortment. An additional benefit of this approach is that we were able to quantify HA-NA pairing strength for individual branches within the phylogenies; given that we demonstrate the cost indicates genetic novelty, this measure may be used as a triage mechanism to identify representative viruses for characterization within risk assessment pipelines or for studies on virus biology [Nguyen et al., 2025, Yamaji et al., 2025].

**Fig. 4.**
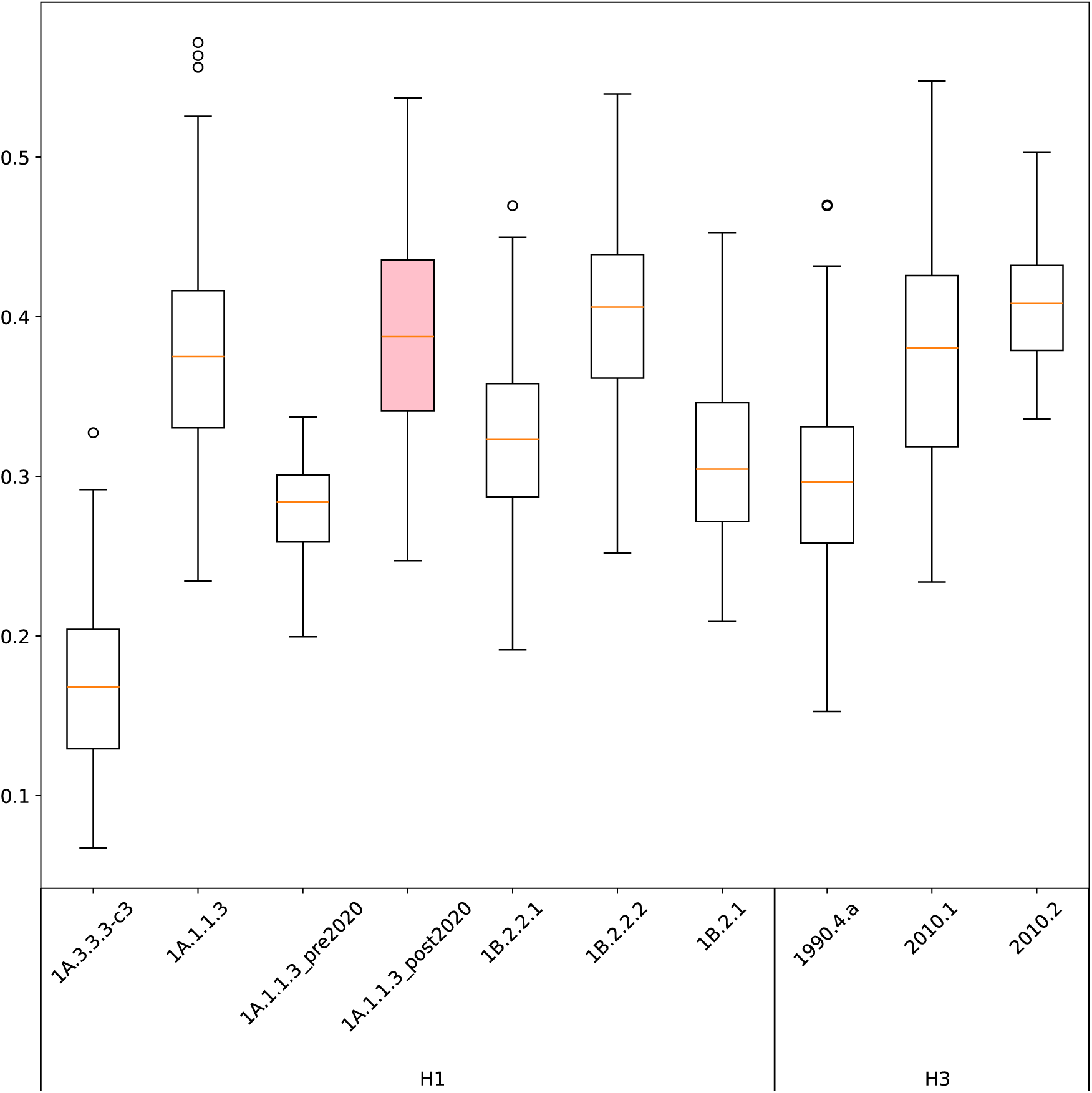
Box and whisker plot of the ACS cost from the hemagglutinin (HA) gene segment phylogenies to the neuraminidase (NA) gene segment phylogenies for H1 and H3 subtype influenza A viruses in swine. The highlighted box represents the post-2020 1A.1.1.3 HA and NA phylogenies which showed an increased ACS cost due to novel gene pairings. The clades are sorted in ascending order of their time of introduction into swine and separated into H1 and H3 subtypes. We observed a near linear increase in the ACS costs that was correlated with the time of introduction and duration of circulation in swine.

## Discussion

Coevolutionary signal can be measured as the topological discordance between the underlying phylogenies [Avino et al., 2019a,b]. Phylogenetic distances quantify this discordance by either utilizing an existing evolutionary model [Górecki and Tiuryn, 2006] or converting the trees to an equivalent mathematical representation [Robinson and Foulds, 1981, Lin et al., 2011]. However, most phylogenetic distances are unable to accurately represent the underlying asymmetry in the phylogenies, for example, in the case of gene tree to species tree comparisons [Degnan and Rosenberg, 2006, Smith et al., 2015] or in the case where there is sampling bias [Bush et al., 2000]. To enable such asymmetric comparisons, phylogenetic costs which relax the requirements of symmetry required by phylogenetic distances were introduced. In this study, we evaluated the performance of two asymmetric phylogenetic costs, the ACA and ACS costs, as indicators of coevolutionary signal.

Identifying rates of cross-species transmission and determining factors that affect viral coevolution or divergence can facilitate our understanding of viral evolution [Olival et al., 2017, Emerman and Malik, 2010]. A coevolutionary study by Geoghegan et al [Geoghegan et al., 2017] quantified family-level interspecies transmission, but was unable to show strong support for variation between the families using an unrooted version of the RF distance (the nPH85 distance). This phylogenetic study contrasted prior research that demonstrated the rate of interspecies transmission and host switching exhibited significant variation based on viral and host families [Olival et al., 2017, Kaján et al., 2020]. Our re-analysis of the Geoghegan et al [Geoghegan et al., 2017] data was able to qualitatively replicate their findings, and demonstrate that the ACA and ACS costs could detect major evolutionary events such as interspecies transmission within the topology of the phylogenies. Our analysis also revealed that the ACA and ACS costs have finer-resolution than Robinson-Foulds-based distances, demonstrated by a broad range of scores that quantified variation between the degree of host-switching across viral families. Our analyses showed that the frequency of interspecies transmission varied significantly across virus families and can be applied to prioritize surveillance and characterization efforts [Carroll et al., 2018].

Virus genes may coevolve resulting in preferential gene pairings [Escalera-Zamudio et al., 2020, Neverov et al., 2015] that can be detected in phylogenetic tree topologies. This is likely the result of biological functions such as RNA-RNA interactions [Jones et al., 2021], or functional restrictions that require a balance between different genes and their encoded protein activity. For example, in influenza A virus, HA-NA pairings are regulated by vRNA segment interactions [Jones et al., 2021] and the two genes have higher fitness when they function in balance [Gaymard et al., 2016]. Our analysis suggests that a similar process occurs in Bluetongue virus (BTV), evidenced by segments 2 and 6 displaying a much lower ACS cost, and by extension, a stronger coevolutionary signal between their respective gene phylogenies. This is likely the result of the roles segment 2 and 6 play in cell entry and exit mechanisms [Roy, 2017, Forzan et al., 2007]. That this signal is maintained as the virus is transmitted between vertebrate hosts and arthropod vectors suggests that there are similar selective pressures within both hosts. Interestingly, both phylogenies also diverge in a similar manner from other gene segments, which we hypothesize reflects a relaxation of the preferential gene pairings as these genes have different biological functions and likely different selective pressures within the vertebrate and invertebrate hosts.

Our data also demonstrated a much higher ACS cost from and to the segment 10 phylogeny from all other gene segment phylogenies. This is plausible as the segment 10 gene has a relatively short evolutionary history in the transmission and evolution of BTV in the US [Alkhamis et al., 2020]. Moreover, the segment 10 gene encodes for the NS3, NS3a and NS5 proteins [Rodríguez-Martín et al., 2021, Mohd Jaafar et al., 2023], which are involved in viral egress [Han and Harty, 2004] and supporting viral replication [Mohd Jaafar et al., 2023], and hence experience stronger selective pressure further contributing to the divergence observed in the ACS cost. Our data also shows lower ACS costs between Seg1, Seg3 and Seg9 phylogenies. This supports previous analyses on European strains of BTV [Nomikou et al., 2015] that showed an association between the viral proteins encoded by segments 1, 3 and 9 as these proteins form the viral replication complex. The same analyses also found associations between segment 2 and 6 evolutionary histories, while confirming the independence of segment 10 evolution.

Our analysis of influenza A virus in swine revealed that the surface proteins had preferential pairings [Hufnagel et al., 2023, Zeller et al., 2021]. In addition, though it was weak, we detected a trend towards stronger coevolutionary dynamics in viruses that had been circulating in swine for longer periods. Intuitively, newer HA clades that have novel HA-NA pairings should have lower coevolutionary signal and, by extension, more topological discordance. Considering that HA-NA pairings are the main drivers for transmission of IAV [Neumann et al., 2009, Neverov et al., 2015], we also expect HA clades that have been in pigs for longer to display lower discordance as the HA-NA pairings should stabilize over time. Our data shows that the ACS cost identifies these patterns, showing a weak but near-linear increase for the H1 and H3 clades from older to newer clades. The exception to this pattern was the H1 1A.1.1.3 clade, likely due to multiple reassortment events within the clade with new HA-NA gene pairings. Separating the 1A.1.1.3 HA clade into phylogenies reflecting the pre- and post-reassortment gene sequences supports this proposition, with the post-reassortment sequences exhibiting a higher ACS cost due to recent reassortment events [Janzen et al., 2025]. The 1B.2.2.2 phylogenies also showed a higher ACS cost that deviated from the general trend, though in this case, the pattern is likely an artifact from randomly downsampling the trees to 50 taxa. Calculating the ACS cost for the complete 1B.2.2.2 trees showed an ACS cost value that fit the linear trend over time. Our approach sought to minimize bias through random sampling, but in this case, our experimental design influenced the result. There were approximately 767 taxa in the 1A.3.3.3 phylogeny while the 1B.2.2.2 clade contained only 273 taxa; this difference, combined with the Gaussian distribution exhibited by the ACS cost [Wagle et al., 2024], allowed the 1A.3.3.3 phylogeny to have a larger range for the ACS cost and consequently a higher probability to capture a meaningful result over randomly sampling taxa.

A major result that emerged from this analysis was that the ACS cost can be applied to detect novel HA-NA gene pairings. Reassortment of IAV in swine typically reflects unique interspecies transmission events and mixing of those viruses in swine hosts [Markin et al., 2023]. All recent IAV pandemics are attributable to a reassortant strain with novel gene pairings [Neumann et al., 2009]. Hence, quantifying the stability of HA-NA pairings in a clade and identifying novel HA-NA pairings is crucial for pandemic risk assessment [Yamaji et al., 2025, Markin et al., 2025].

Coevolutionary studies may reflect asymmetry in the input data and phylogenies [Bush et al., 2000, Degnan and Rosenberg, 2006, Smith et al., 2015], and the degree of asymmetry between two phylogenies is often unknown or indeterminate. In this study, we demonstrated the applicability of asymmetric cluster-based costs to compare phylogenies, allowing a more robust indicator for coevolutionary signal even when the input data is potentially noisy and the phylogenies have an asymmetric relationship. The coevolutionary signals identified by these costs provided insight into viral evolution, such as identifying coevolutionary pairings between virus families and hosts, characterizing gene pairings in viral segments to detect coevolutionary dynamics as viruses are transmitted between vertebrate and invertebrate animals, and detecting novel gene pairings. Future research should focus on incorporating additional measures of topological dissimilarity such as branch lengths, comparison between other phylogenetic structures, experimental characterization of taxa across the breadth of the score measure to determine its functional relevance, and modeling approaches that are able to enrich the phylogenetic cost with additional taxa-level information such as frequency of interactions between taxa and the probability for those interactions to be antagonistic or mutualistic.

## Acknowledgements

The authors gratefully acknowledge pork producers, swine veterinarians, and laboratories for participating in the USDA Influenza A Virus in Swine Surveillance System and publicly sharing sequences in NCBI GenBank. This project was funded in part by the United States Department of Agriculture (USDA), Agricultural Research Service (ARS project numbers 3022-32000-018-017-S, 5030-32000-231-103-A, 5030-32000-231-095-S, 5030-32000-231-000-D) and with federal funds from the National Institute of Allergy and Infectious Diseases, National Institutes of Health, Department of Health and Human Services (contract No. 75N93021C00015) and the SCINet project and the AI Center of Excellence of the USDA ARS (ARS project numbers 0201-88888-003-000D and 0201-88888-002-000D). The funding sources had no role in study design, data collection, and interpretation, or the decision to submit the work for publication. USDA is an equal opportunity provider and employer.

## Data Availability Statement

All ACA and ACS costs were computed using the cluster affinity software package available at https://github.com/swagle8987/cluster_affinity. Phylogenetic trees used in the Bluetongue virus and influenza A virus analysis are available on Dryad.

